# A gated relaxation oscillator controls morphogenetic movements in bacteria

**DOI:** 10.1101/137695

**Authors:** Mathilde Guzzo, Seán M. Murray, Eugénie Martineau, Sébastien Lhospice, Grégory Baronian, Laetitia My, Yong Zhang, Leon Espinosa, Renaud Vincentelli, Benjamin P. Bratton, Joshua W. Shaevitz, Virginie Molle, Martin Howard, Tâm Mignot

## Abstract

Dynamic control of cell polarity is of critical importance for many aspects of cellular development and motility. In *Myxococcus xanthus*, a G-protein and its cognate GTPase-activating protein establish a polarity axis that defines the direction of movement of the cell and which can be rapidly inverted by the Frz chemosensory system. Although vital for collective cell behaviours, how Frz triggers this switch has remained unknown. Here, we use genetics, imaging and mathematical modelling to show that Frz controls polarity reversals via a gated relaxation oscillator. FrzX, which we newly identify as the primary Frz output, provides the gating and thus acts as the trigger for reversals. Slow relocalisation of the polarity protein RomR then creates a refractory period during which another switch cannot be triggered. A secondary Frz output, FrzZ, decreases this delay allowing rapid reversals when required. This architecture thus results in a highly tunable switch that allows a wide range of motility responses.

## Introduction

Periodic dynamics are pervasive in biology with examples ranging from circadian rhythms to brain activity and from the cell cycle to multicellular development^1^. Study of rhythmic processes in microbes has often been particularly insightful due to the simpler nature of the system dynamics. Classic examples include the *Escherichia coli* Min system regulating cell division positioning and the circadian Kai oscillator in cyanobacteria^2,3^.

Rhythmic processes can be particularly important in microbes for the regulation of cell motility. In the bacterium *Myxococcus xanthus*, back and forth movements (reversals) are not only important for the motile dynamics of isolated cells but are also critical for the regulation of multicellular behaviours, fruiting body formation and rippling during the invasion and consumption of prey cell colonies^4^. The function of periodic reversals is especially evident during rippling, the accordion-like wave movements of large cell groups, where each cell of a wave reverses upon collision with a cell of an opposite incoming wave^5^. This large-scale order has been proposed to emerge from the synchronisation of an intracellular compass (called Frz) by cell-cell contacts^5,6^. However, in other contexts *M. xanthus* reversals can be aperiodic^4^, and thus the underlying mechanism that generates reversals must be able to create richer dynamics than simple oscillations. In this study, we use an interdisciplinary approach combining genetics, live imaging and mathematical modelling, to elucidate this underlying mechanism, dissecting how Frz controls dynamic polarity in individual cells. Such an interdisciplinary strategy is essential to properly dissect how the individual molecular components fit together to generate coherent dynamics and rapid switching.

*Myxococcus* cells employ two motility systems depending on the context, both of which are assembled at the leading cell pole. The S-motility complex functions within large cell groups and consists of a Type-IV pilus that deploys to pull the cell forward^7^ (Figure 1A), while the A-motility complex traffics toward the lagging cell pole, propelling the cells as it becomes adhered at so-called bacterial focal adhesions^8^ (Figure 1A). Thus at the molecular level, cell reversals are provoked by the rapid activation of the motility complexes at the opposite cell pole (Figure 1A). The genetic control of cell reversals involves a chimeric circuit composed of bacterial Che-like proteins and a eukaryotic Ras-like regulation system (Figure 1A). Spatial activation of A-and S-motility depends on a single regulator, MglA, a G-protein of the Ras superfamily, which binds to the leading cell pole in its active GTP-bound form and recruits key proteins of each motility system^9–12^ (Figure 1A). The activity of MglA is regulated by MglB, a GTPase Activating Protein (GAP) that binds to MglA in a 2:1 stoichiometry to activate the transition from polar-localised MglA-GTP to cytoplasmic MglA-GDP^13^. Since MglB localises to the lagging cell pole, its GAP activity is spatially regulated, blocking access of MglA to the lagging pole. This MglAB polarity axis can be inverted on a timescale of 30-60 seconds, leading to cell reversals^9,12^.

**Figure 1.**
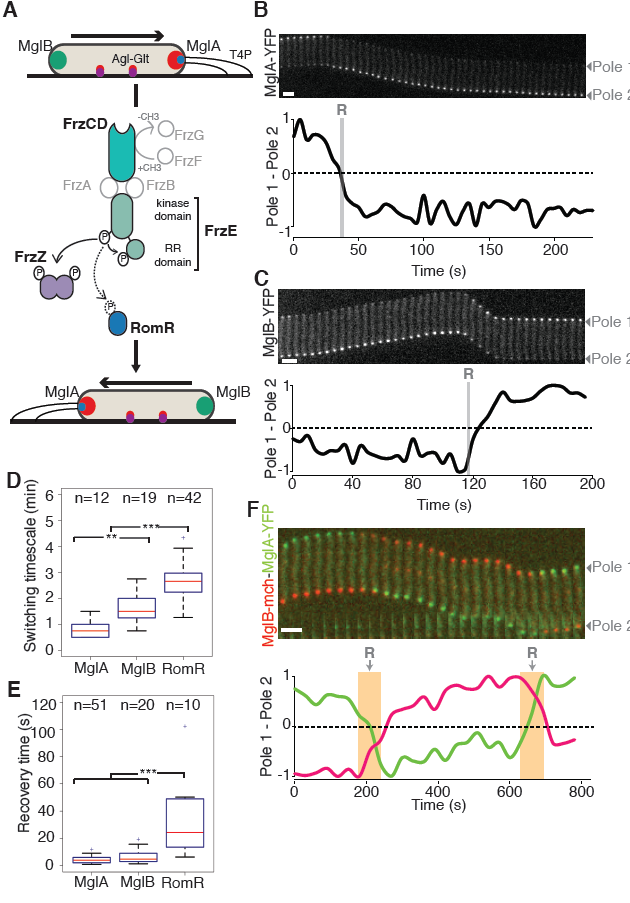
Pole-to-pole dynamics of the polarity proteins. **(A)** Genetic structure and function of the Frz pathway, based on previously published work. **(B)** Top: Timelapse images of pole-to-pole dynamics of MglA-YFP (5s time frames) in a single reversing WT cell. Bottom: difference in intensity of MglA-YFP between the two poles normalized to maximum absolute value of difference shown as a function of time. R: Reversals, detected by a clear directional change of the cell. Scale bar = 2 μm. **(C)** As in (B), but for MglB-YFP in a single reversing WT cell. **(D)** Boxplots of the switching timescale distributions for MglA-YFP, MglB-YFP and RomR-GFP. The lower and upper boundaries of the boxes correspond to the 25% and 75% percentiles, respectively. The median is shown as a red line and the whiskers represent the 10% and 90% percentiles. ** p-value<0.01, *** p-value<0.001 **(E)** Boxplots of the recovery timescales for MglA-YFP, MglB-YFP and RomR-GFP as determined by FRAP. Boxplots read as in (D). *** p-value>0.001 **(F)** Top: Time-lapse images of pole-to-pole dynamics of MglA-YFP and MglB-mCh (5 s time frames) in a single reversing cell. Bottom: differences in intensities of MglA-YFP (green) and MglB-mCh (red) between the two poles, normalized to the respective maximum absolute value of the difference, shown as a function of time. Note that MglA-YFP switches first and that both proteins co-localise for a short period immediately before reversals. Scale bar = 2 μm.

The mechanism by which polarity is switched is, however, still not understood despite intense investigation. Upstream, the switch is controlled by the Frz chemosensory-like system, which forms signalling arrays at discrete positions along the bacterial nucleoid^14^, and connects to the CheA-type kinase FrzE. Following activation (by as yet unknown physiological signals, possibly cell-cell contacts), FrzE dimers have been proposed to phosphorylate several downstream Response Regulator (RR) domain proteins: the cognate FrzE RR domain FrzE^RR^, the tandem RR protein FrzZ and the RR domain of RomR (RomR^RR^)^4,15–17,14^ (Figure 1A). The respective contribution of each RR domain has been RR investigated by genetic analysis, suggesting that the central output of the pathway is RomR^RR^ ^4,18^ FrzE^RR^ and FrzZ act as accessory domains that impact the signal flow negatively and positively, respectively (Figure 1A). Specifically, FrzE^RR^ acts as a phosphate sink preventing noisy activation of the system at low stimulation levels^4,14^. On the contrary, the non-essential FrzZ protein amplifies the system efficiency by an as-yet undetermined mechanism^4^ (Figure 1A). However, contrary to FrzE^RR^ and FrzZ^15,16^, it has not been shown experimentally that RomR^RR^ is a direct substrate of the FrzE kinase. In motile cells, RomR interacts with MglA-GTP and is essential for MglA’s polar localisation^19,20^. However, RomR also interacts with MglB at the lagging cell pole^19,20^. While the exact function of the RomR localisation pattern is not understood, RomR is known to be essential for cell reversals^4^. Therefore, direct phosphorylation of RomR has been proposed to connect Frz signalling to the MglAB polarity complex.

In this study, we identify for the first time the link between Frz and the polarity proteins. We find that Frz, via its newly discovered primary output FrzX, acts as the trigger for a new type of biological oscillator: a gated relaxation oscillator. Here, we use the terminology of electronic circuits, where ‘gated’ refers to the requirement for an input signal, in this case FrzX, to trigger reversals^21^. The oscillator itself consist of the polarity proteins MglA, MglB and RomR where RomR sets the concentration of MglA both at the leading and lagging poles. The slow dynamics of RomR further introduces a refractory period (relaxation time) immediately after a switch during which time another reversal cannot be triggered. However, Frz is able to overcome this minimum period to achieve rapid reversals when required via the action of its secondary output FrzZ, explaining the previously observed accessory function of FrzZ. This unique system design allows a wide range of responses to variations in incoming signals.

## Results

### MglA and MglB rapidly switch their polar localisation when cells reverse

Polar switching only occurs in the presence of all three polarity proteins MglA, MglB and RomR, indicating that they are all part of the same oscillatory circuit. However, how the phosphorylation of RomR could control the switch is not known^4,18^. To investigate this mechanism, we first determined the sequence of events that leads to switching, performing high-time resolution fluorescence microscopy using YFP fusions to MglA and MglB^12^ (Figure 1B,C), followed by quantitative image analysis (see Supplementary Information). In this assay, both proteins re-localised at the time of reversals as previously described^12^, with switching timescales (i.e. the time necessary for the entire protein fluorescent cluster to switch pole, see Figure S1A, Methods) of 45 ± 20 s (n=12) for MglA-YFP and 90 ± 30 s (n=19) for MglB-YFP (Figure 1D). Fluorescence Recovery After Photobleaching (FRAP) revealed the polar dissociation of MglA and MglB is rapid with mean recovery times of 4 s (n=51) and 6 s (n=20) respectively (Figure 1E, Figure S1B, Methods). The MglA and MglB recovery times were not substantially affected in *frz* mutants (Figure S1C,D), suggesting that Frz signalling does not directly modulate the affinity of MglA and MglB for the poles. Analysis of a dual colour strain expressing both MglA-YFP and MglB-mCherry^12^ revealed that when cells reverse, both proteins co-localise during a short time window and MglA is the first protein to re-localise during a given reversal (observed in 10 out of n=10 events, Figure 1F). This suggests that overcoming the repelling action of MglB triggers a reversal event (Figure 1F)

### A three-protein relaxation oscillator model of the reversal cycle

We next turned to mathematical modelling to investigate how interactions between MglA, MglB and RomR could generate the oscillations. Our initial mathematical model incorporated observations from the previous section, as well as those previously published (Supplementary Information). In particular, we assumed that MglA and MglB exert bidirectional antagonistic effects such that each protein excludes the other protein from the pole, with local concentrations determining which protein is dominant over the other at a given pole (Figure 2A,B Supplementary information). This could be explained if immediately after a reversal, MglB efficiently inhibits polar MglA-GTP by stimulating GTP hydrolysis^9,12^ returning MglA to the cytoplasm in the GDP form. However, as the cell gets closer to the next reversal, MglB also sequesters RomR progressively, and this interaction in turn increases the recruitment rate of MglA-GTP^19,20^. Eventually MglB inhibition of MglA becomes saturated when RomR accumulates to sufficient levels (Figure 2B). At this point, excess MglA-GTP could effectively replace MglB if the MglA-MglB interaction detaches MglB from the pole (Supplementary Information). MglB would thus be displaced to the other pole where MglA-GTP levels are significantly lower, allowing the process to begin again. This process would be greatly facilitated if MglB has an affinity for itself (through a cooperative self-interaction). Consistent with this hypothesis, structural studies indicate that MglB can form tetramers^13^. Furthermore, MglB is monopolar in an *mglA romR* mutant^20^, a localisation pattern that could be explained if MglB cooperatively forms polar oligomers similar to, for example, the hub protein PopZ^22,23^. We therefore incorporated into the model an affinity of MglB for itself. The equations describing the interactions in Figure 2A (solid and non-solid arrows) are presented in Figure 2C.

**Figure 2.**
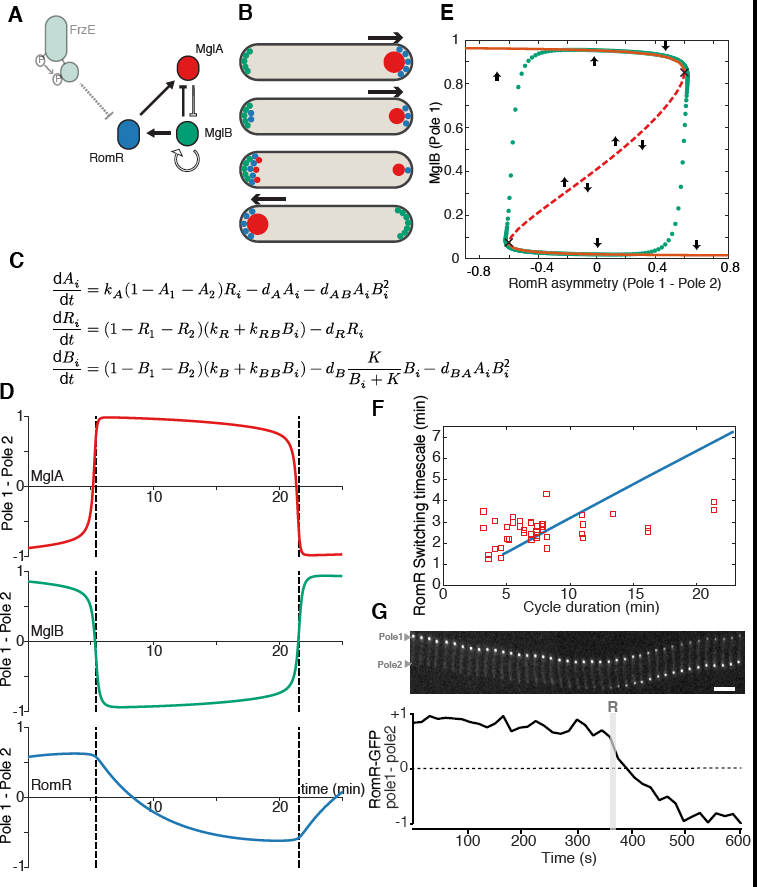
A three-protein relaxation oscillator model of polarity reversals. **(A)** Schematic of the polarity module. Solid arrows indicate interactions supported byexisting data. Hypothetical interactions indicated by non-solid arrows. Blunt-end arrows refer to negative effects on polar localisation at the same pole. The Frz chemosensory system is proposed to modulate the timescale of RomR dynamics, possibly by direct phosphorylation. **(B)** Schematic localisation of polarity proteins with respect to direction of movement inferred from the interactions proposed in (A). Proteins colored as in (A). **(C)** The model consists of six ordinary differential equations describing the relative polar levels of MglA (A_i_), RomR *(R)* and MglB (B_i_) at each pole, i=1,2. Binding and unbinding rates are indicated by *k* and *d*, respectively. See Supplementary Information for further details and parameter values. **(D)** Profiles produced by equations in (C), showing difference in levels between the two poles. Dashed lines indicate the time of parity of the MglB levels. **(E)** Bifurcation diagram showing fixed points of *Bi* as a function of RomR asymmetry *(R_1_* -*R_2_)* (orange line; the stable branch is solid, unstable is dashed, as also indicated by arrows). The saddle bifurcations are indicated by crosses. Overlayed in green is the solution shown in (D). The solutions proceeds clockwise and the data points are evenly spaced in time. **(F)** Correlation between the RomR switching timescale and the cycle duration as predicted by the model (blue line), and as obtained experimentally (red squares). Model data was obtained by measuring the cycle duration and switching timescale as for experimental profiles. Different cycle durations were obtained by scaling all RomR rates (binding and unbinding) by the same factor. The model predicts that the RomR switching timescale should be strongly correlated to the cycle duration. However, the observed correlation is weak: Pearson’s correlation coefficient of the experimental data is r=0.45 (p-value p=0.003, calculated using Student’s t-distribution). Experimental data is from a mix of unstimulated and stimulated (0.03% IAA) WT cells. **(G)** Example RomR-GFP profile as in Fig. 1B showing both the slow accumulation of RomR-GFP at the lagging pole and the stability of its levels before a reversal. Scale bar = 2 μm.

We solved this system of equations numerically and found that the system could indeed exhibit oscillations, with the MglA and MglB profiles qualitatively similar to the experimental curves (Figure 2D). Specifically, the oscillations were driven by RomR binding to MglB and then recruiting MglA, with switching provoked when RomR reaches a critical threshold at the lagging pole. This driving could be seen explicitly by manually varying the asymmetry in polar RomR levels and observing the effect on polar MglB localization (Figure 2E). We also observed that oscillations only took place if the dynamics of RomR occurred on a timescale longer than that for MglA and MglB. We further found that, in this case, the slower RomR timescale set the cycle duration (defined as the time between two successive polarity switching events) (Figure 2F). These properties are characteristic of a relaxation oscillator, in which oscillations are due to, and set by, a component of the system with dynamics much slower than the others. Indeed, from simulations we found a linear relationship between the timescale of RomR dynamics and the cycle duration (Figure 2F). Thus, if phosphorylation of RomR by the FrzE kinase (Figure 2A) alters the (slow) timescale of RomR dynamics, then the model suggests that this regulation would specify the cycle duration (Figure 2D,F).

### RomR dynamics do not time the reversal cycle

We next tested if our experimental observations of RomR are indeed consistent with the slow dynamics predicted by the model. We found that RomR switching indeed occurred on longer timescales than MglA and MglB (mean 160 s, Figure 1D) with a mean FRAP recovery time of 28s, slower than that measured for both MglA and MglB (Figure 1E). Unexpectedly however, in WT cells RomR switching dynamics were not correlated to reversals and RomR-GFP accumulated stably at the lagging pole before the cell reversed (Figure 2G). In fact, the timescale of RomR switching was largely constant and did not vary significantly as a function of the cycle duration (Figure 2F). These results suggest that switching cannot be solely regulated by the slow dynamics of RomR, a conclusion that questions whether RomR is indeed a substrate of the FrzE kinase.

We therefore tested if the receiver domain of RomR (RomR^RR^, Figure S2A) is phosphorylated by FrzE *in vitro*. Consistent with published results^15^, purified FrzE phosphorylated FrzZ very efficiently, achieving 100% transfer less than 2 min after the proteins were mixed (Figure 3A). However, under the same conditions, neither full length RomR nor RomR^RR^ were significantly phosphorylated even after 8 min incubation (Figure 3A, Figure S2B).

**Figure 3.**
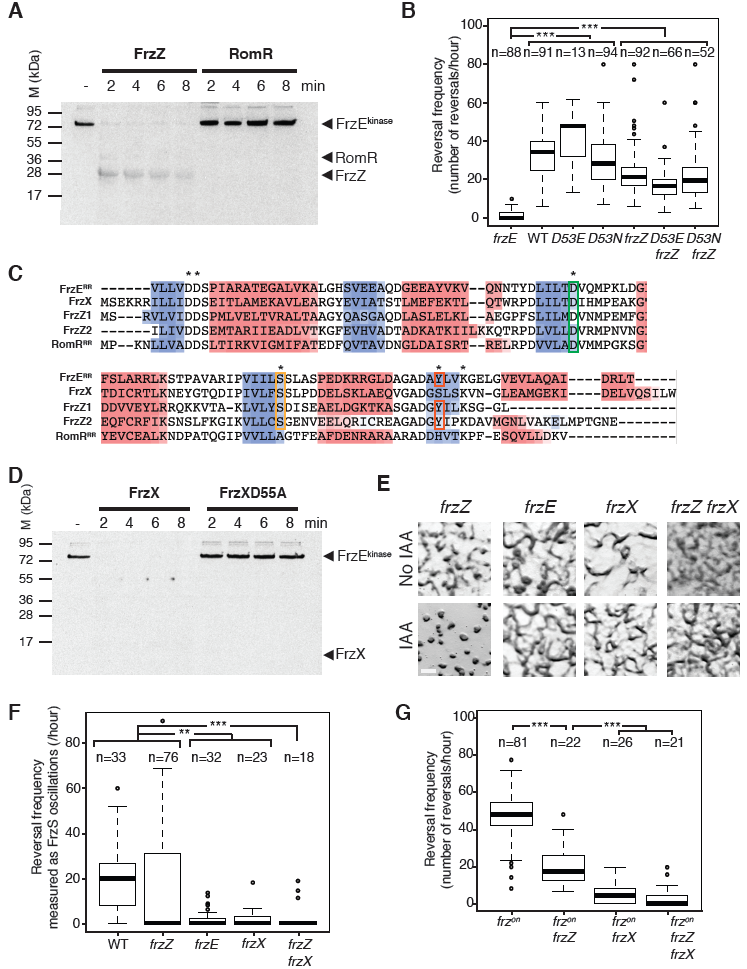
FrzX not RomR is the major FrzE kinase substrate used to control the reversal cycle. **(A)** RomR is not phosphorylated by the FrzE kinase *in vitro*. Autoradiogram of P^32^-labelled FrzE in the presence of FrzZ or RomR. Note that P^32^ is readily transferred to FrzZ but not to RomR. **(B)** Single-cell S-motility reversal frequency of WT and selected mutants on carboxymethylcellulose (see Methods) in the presence of 0.15% IAA. Shown are boxplots of the measured reversal frequency monitored as the number of directional changes per hour (see Methods) of isolated cells (tracked for at least 10 min) for each strain. Boxplots read as in Figure 1D. D53E: *romR^D53E^*, D53N: *romR^D53N^*. Statistics: Wilcoxon test (n<40) for *rom^53E^* and student test (t-test, n>40) for all the other strains. *** p-value<0.0001. Note that the WT, *frzE* and *frzZ* controls were published previously^4^ (see Methods). **(C)** Multiple sequence alignment of Frz-associated RR proteins. Sequence alignment was generated using MAFFT and secondary structures were predicted using Ali2D (a-helices: residues highlighted in red. P-sheets: residues highlighted in blue). Stars indicate the phosphorylatable aspartate residues (D) and critical residues important for signal transduction Serine (S) and Tyrosine (Y). **(D)** FrzX is directly phosphorylated by the FrzE kinase *in vitro*. Direct phosphorylation is observed through the rapid disappearance of P^32^-labelled FrzE in the presence of FrzX. A FrzX^D55A^ mutant does not affect P^32^-FrzE indicating that this effect is due to a phosphotransfer event. **(E)** FrzX is a central output of the Frz pathway. Developmental phenotypic assays showing the typical “frizzy” phenotype of *frzX* mutants. Note that IAA rescues the development of a *frzZ* mutant but not that of either *frzE* or *frzX* mutants, indicating that FrzX is central for the regulation. Scale bar: 1mm. **(F)** FrzX is essential for reversals. Single-cell S-motility reversal frequency on carboxymethylcellulose in the presence of 0.1% IAA. Shown are boxplots of the measured reversal frequency monitored as the number of FrzS-YFP oscillations per hour (see Methods) of isolated cells (tracked for at least 10 min) for each strain. The boxplots read as in Figure 1D. Statistics: Wilcoxon test (n<40) for WT, *frzE, frzX* and *frzZ frzX* and student test (t-test, n>40) for *frzZ*. ** p-value<0.01, *** p-value<0.0001. Note that the WT, *frzE* and *frzZ*controls were published previously^4^ (see Methods). **(G)** FrzX but not FrzZ is essential for reversals in Frz-hyper signaling mutants *(frz^on^)*.Single-cell S-motility reversal frequency of *frz^on^* and selected mutants on carboxymethylcellulose (see Methods) without IAA. Shown are boxplots of the measured reversal frequency monitored as the number of directional changes per hour (see Methods) of isolated cells (tracked for at least 10 min) for each strain. Boxplots read as in Figure 1D. Statistics: Wilcoxon test. *** p-value<0.0001. Note that the *frz^on^* and *frz^on^ frzZ* controls were published previously^4^ (see Methods).

Since we obtained no evidence for RomR phosphorylation *in vitro*, we revisited the impact of point mutations on the conserved D53 residue of RomR^RR^ *in* vivo^18^ by substituting *romR* with a *romRP^D53E^* (phospho-mimicking mutation) or *romR^D53N^* (phospho-ablative mutation). To quantitatively measure the impact of the point mutations on the reversal frequency, we scored single cell reversals of both mutants as compared to the WT in a microfluidic device where the level of Frz activation can be controlled by Isoamylalcohol (IAA), a known Frz activator^4^ (see methods). Surprisingly, the *romR^D53E^* and the *romR^D53N^* mutants showed no detectable defect and had a reversal frequency comparable to the WT (Figure 3B). In addition, the *romR^D53E^ frzZ* and *romR^D53N^ frzZ* double mutants showed reversals similar to the *frzZ* mutant^4^ (Figure 3B), while it is known that the *romR frzZ* double mutant does not reverse, similar to the *romR* mutant^4^. Thus, the presence of RomR, but not its phosphorylation, is required for reversals. Consistently, the RomR point mutant strains showed no developmental defects: both the *romR^D53E^* and *romR^D53N^* point mutants formed fruiting bodies on hard surfaces, showing that the phosphorylation of RomR is not critical for multicellular development (Figure S2C). We therefore conclude that RomR is required for reversals, presumably because it is needed for the polar localisation of MglA but its phosphorylation is not involved in the control of the switch. Furthermore, since FrzE can still trigger reversals when the *frzZ* gene is deleted and/or when the RomR^RR^ receiver domain is mutated, these results implicate an as yet unidentified response regulator in the reversal process.

### Identification of a new target of the FrzE kinase

To identify the missing regulator, we reasoned that it should share structural similarities with FrzZ, which is directly phosphorylated by FrzE. The search of close FrzZ homologs identified the product of the MXAN_5688 gene as a predicted single domain response regulator protein with high sequence homologies to the FrzZ1 domain (Figure 3C). In addition, a homolog of MXAN_5688 is encoded in proximity to a frz-type operon in *Anaeromyxobacter* species, which contains both a FrzZ and an MXAN_5688 homolog (Figure S3A). To test if the protein encoded by the MXAN_5688 locus is a direct target of FrzE, we purified the protein (named FrzX) and tested its phosphorylation *in vitro*. Co-incubation between the FrzE kinase domain and FrzX led to the complete disappearance of phosphorylated FrzE even after 30 sec of co-incubation, suggesting that phosphotransfer had occurred (Figure 3D). However, no band corresponding to phosphorylated FrzX could be detected. The phosphorylated state of RR domains with high auto-phosphatase activity can be difficult to capture on SDS-PAGE gels^24,25^. Therefore, to test whether FrzE indeed phosphorylates FrzX, we repeated the phosphotransfer assay in the presence of a FrzX variant carrying a non-phosphorylatable D55A mutation, where D55 is the predicted phosphorylatable Asp (Figure 3C,D). Under these conditions, FrzE remained phosphorylated even after 8 min co-incubation (Figure 3D), suggesting that FrzX is indeed a direct target of the FrzE kinase.

### FrzX is the missing Frz Response Regulator protein

Similar to a frzE-null mutant, a *frzX-null* mutant formed typical “frizzy” tangled filaments instead of fruiting bodies on developmental agar plates (Figure 3E). The deletion of *frzX* could be complemented with a *frzX* allele but not with a *fizX^D55A^* allele, showing that the phosphorylation of FrzX is essential for its function (Figure S3B,C). FrzX is central to Frz signalling because neither a *frzX* mutant nor a *frzX frzZ* double mutant were rescued by the addition of IAA (Figure 3E). By contrast, IAA addition can rescue the absence of FrzZ, implying that, despite its signal amplification properties, FrzZ is a less critical component of the reversal apparatus^4^ (Figure 3E).

To show directly that FrzX acts in the regulation of reversals, we measured the reversal frequency of the *frzX* mutant in a single cell assay that measures IAA and Frz-dependent protein switching directly^4^ (FrzS-YFP). This methodology can thus unambiguously distinguish *bona fide* reversals from other movements such as stick-slip motions (these motions occur at a low level adding noise to standard reversal scoring assays^4^, see Methods). As expected, the *frzX* mutant failed to reverse, similar to a *frzE* kinase mutant (Figure 3F), but dissimilar to the *frzZ* mutant which does still reverse in this assay, albeit at a lower frequency, in the presence of IAA (Figure 3F). These results again place FrzX in a more central position in the reversal mechanism as compared to FrzZ.

To prove this central function of FrzX in a definitive manner, we tested its contribution in a strain with a hyper-signaling state of the FrzE kinase (termed the *frz^on^* mutation^4,26^, where the cells hyper-reverse due to constitutive activation by the Frz receptor (FrzCD). These reversals can be detected in our single cell assay in the absence of IAA (Figure 3G). A *frz^on^frzZ* mutant still reversed, although again at a lower frequency since FrzZ is needed for optimal signalling activity^4^. In contrast, both a *frz^on^ frzX* and a *frz^on^ frzX frzZ* mutant failed to reverse (Figure 3G). We conclude that FrzX acts genetically downstream from FrzZ and, unlike FrzZ, is absolutely required for the transmission of FrzE signals to the downstream polarity proteins.

### The phosphorylated form of FrzX localises to the lagging cell pole in an MglB-dependent manner

To understand how FrzX provokes cellular reversals downstream from the FrzE kinase, we first tested how deleting *frzX* affects the localisation of MglA-YFP, MglB-YFP and RomR-GFP. In a *frzX* mutant, these proteins were correctly localised at their respective poles (Figure S4A) but they did not switch poles, consistent with FrzX being essential for reversals. Thus, FrzX is only required for the dynamic switching of the polarity proteins.

We next constructed a functional N-terminal GFP-FrzX fusion (Figure S3D) to investigate where FrzX localises. We found that GFP-FrzX was present diffusely in the cytoplasm when Frz-signalling is at low level (Figure 4A). However, the addition of IAA induced the appearance of a clear polar focus (Figure 4A). This polar localisation was fully abrogated in a *frzE* mutant in the presence of IAA (Figure 4A). These results suggest that phosphorylation of FrzX by FrzE is required for polar localisation.

**Figure 4.**
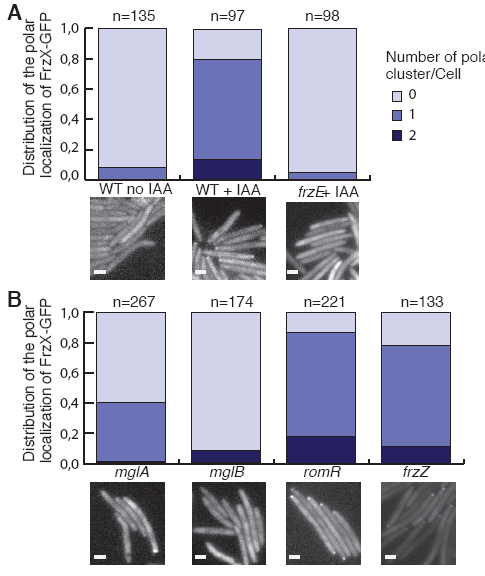
FrzX localises to the cell pole in phosphorylation-and MglB-dependent manners. **(A)** FrzX localises to the cell pole in a phosphorylation-dependent manner. Polar localisation patterns in the presence of low (WT no IAA, *frzE* + IAA) and high (WT + IAA) Frz-signaling activity. Scale bar = 2 μm. **(B)** FrzX localisation depends on MglB and partially on MglA. All tests done in the presence of IAA to increase the frequency of polar GFP-FrzX clusters. The WT (+IAA) control is shown in **(A)**. Scale bar = 2 μm.

To identify a possible FrzX target at the pole, we next investigated the localisation of GFP-FrzX in the polarity protein mutants, *mglA, romR, mglB* and *frzZ*. Except for the *mglB* mutant, GFP-FrzX was polarly localised in all mutants (Figure 4B, although a slight reduction was also observed in the *mglA* mutant), suggesting that either MglB or an MglB-interacting protein is a polar target for FrzX. Finally, consistent with this probable interaction with MglB or an MglB-interacting partner, GFP-FrzX accumulated at the lagging cell pole between reversals (see below).

### RomR and FrzX both act to provoke reversals

To determine how FrzX and RomR act at the lagging cell pole to provoke reversals, we more closely investigated their localisation dynamics during the reversal cycle. Because the polar localisation of FrzX can only be observed at high Frz signalling levels, we performed all the GFP-FrzX localisation assays in the presence of either IAA or a *frz^on^* mutation (similar results were obtained in both conditions). Remarkably, in these Frz-activated cells, reversals coincided with a peak accumulation of GFP-FrzX, suggesting that FrzX triggers cell reversals (Figure 5A,B and Figure S5A).

**Figure 5.**
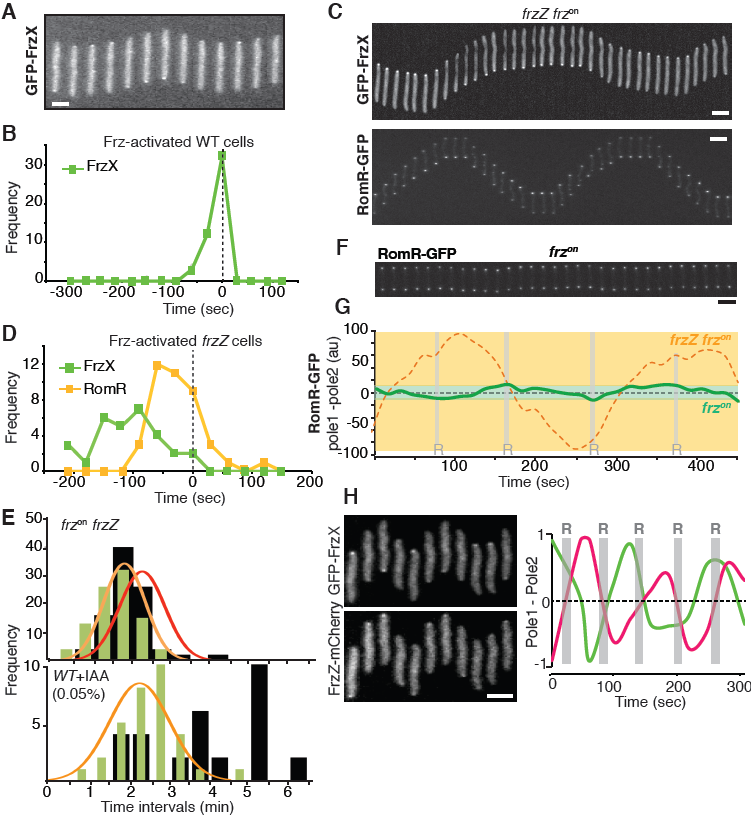
Reversals require the action of both FrzX∼P and RomR at the lagging cell pole. **(A)** Top: Time-lapse images of GFP-FrzX (15 s time frames) in a single cell with IAA-induced high-Frz signalling. Scale bar = 2 μm. See also Figure S5A. **(B)** Reversals correlate with peaks of GFP-FrzX at the lagging cell pole in Frz-activated cells *(frz^on^* or IAA treated). Plot shows frequency of fluorescence peaks (as detected by fluorescence maxima) relative to reversal events. Time=0 indicates the reversal event itself, with negative values representing pre-reversal and positive values representing post-reversal times. **(C)** Time-lapse images (15 s time frames) from single *frz^on^ frzZ* cells expressing GFP-FrzX (top) and RomR-GFP (bottom). Note that GFP-FrzX accumulates stably at the lagging cell pole in this background. Scale bar = 2 m. See Figure S5B,C for IAA treated cells. **(D)** RomR-GFP, but not GFP-FrzX, peaks are correlated with reversals in Frz-activated *(frz^on^*or IAA treated) *frzZ* mutant cells. Otherwise as in (B). **(E)** Reversals are set by the RomR switching time scale in *frz^on^ frzZ* mutant cells. Upper panel: Distribution of RomR-GFP switching times scales (green bars, normal distribution fit in orange) and reversal cycle duration (black bars, normal distribution fit in red) in a *frz^on^frzZ*mutant. Lower panel: same as in upper panel but in WT in the presence of a low concentration of IAA (0.05%, which induces a broad range of reversals). Note that in this case the RomR-GFP switching distribution is little changed and no longer coincides with the distribution of switching timescales. **(F)** Time-lapse images from a single *frz^on^* cell expressing RomR-GFP. Sacle bar = 2 μm **(G)** Difference in RomR-GFP intensities over time between the two poles in a *frz^on^* background (green trace, corresponding reversals are indicated by grey bars) overlaid with the difference in RomR-GFP intensities in a *frz^on^ frzZ* background (orange dotted line, trace corresponding to the cell shown in Figure 5C). Note that for comparison, the pole1-pole2 difference is not normalized to the absolute value of the maximum intensity and thus pole1-pole2 values are expressed in arbitrary units (au). The green area corresponds to the mean amplitude difference (pole1-pole2) between poles at the time of reversal, computed from n = 20 *frz^on^* reversal events, (mean amplitude difference = 12 ± 2). Orange area: same methodology, but for *frz^on^ frzZ* background (from n = 20 reversal events, mean amplitude difference = 98 ± 10). In *frz^on^*, peak asymmetry in polar RomR-GFP intensities is still correlated to reversals, but reversals occur at a 10 fold less amplitude difference than in the *frz^on^ frzZ* background. **(H)** FrzZ and FrzX act simultaneously at opposite cell poles. Left: time-lapse images of the same GFP-FrzX and FrzZ-mCherry hyper-reversing single cell (IAA treated, 15 s time frames). Right: differences in intensities of GFP-FrzX (green) and FrzZ-mCherry (red) between the two poles, normalized to the respective maximum absolute value of the difference, shown as a function of time. R: Reversals, detected by a clear directional change of the cell. Scale bar = 2 μm.

We next tested how GFP-FrzX localises in the absence of FrzZ, again either in the presence of IAA or in a *frz^on^* background. Remarkably, in both conditions, GFP-FrzX still localised to the lagging cell pole but its accumulation no longer coincided with cell reversals (Figure 5C,D and Figure S5B), suggesting that another component is needed for reversals in a *frzZ* mutant. To test if this component could be RomR, we further analyzed RomR-GFP dynamics in Frz-activated cells (Figure 5C and Figure S5C). Indeed, the maximum accumulation of RomR-GFP was strongly correlated with reversals (Figure 5D). Consistent with this, the distribution of the RomR switching timescale coincides with that of the cycle duration in a *frz^on^ frzZ* background (Figure 5E). However, as observed previously, the RomR switching timescale is not significantly correlated with reversals in the wildtype background (Figure 2F, Figure 5E). Taken together the results suggest that both RomR and FrzX function together at the lagging pole to provoke reversals, but which of the two provides the final trigger depends on which is limiting: FrzX in WT but RomR when *frzZ* is missing.

### FrzZ overcomes a refractory period set by the dynamics of RomR

In cells where Frz-signaling is at maximum level, RomR dynamics set a minimum reversal frequency and faster reversals require FrzZ. This function is evident when RomR dynamics are analysed in WT cells in a *frz^on^* background (Figure 5F) and compared to *frz^on^ frzZ* (Figure 5G). In *frz^on^* cells, switching is also correlated to RomR levels at the lagging cell pole but, contrarily to *frz^on^ frzZ* cells, it does not require the entire RomR population to localise to the lagging pole and occurs at a much lower threshold (Figure 5G). This result is confirmed in a dual labelled strain expressing functional GFP-FrzX and RomR-mCherry (RomR-mCh), fusions in which reversals coincide both with peak accumulations of GFP-FrzX and low amplitude changes of RomR-mCh at the lagging cell pole (Figure S5D).

Thus, FrzZ overcomes a limit set by the slow dynamics of RomR, accounting for the previously documented positive effect of FrzZ on the reversal frequency. How FrzZ performs this function is not known. FrzZ localises to the leading cell pole in a FrzE phosphorylation- and MglA-dependent manner^17^, opposite to GFP-FrzX (Figure 5H, Figure S4B). Furthermore, FrzZ and FrzX localise independently to the cell poles (Figure 4B and Figure S4B). In the next section, we will use these results together with mathematical modelling to propose a mechanism for FrzZ function.

### A gated relaxation oscillator model of the polarity switching mechanism, incorporating the functions of FrzX and FrzZ

The above results show that the reversal switch requires the combined action of phosphorylated FrzX and RomR at the lagging pole. By gradually accumulating at the lagging pole via its interaction with MglB, RomR primes the cell for the next reversal. The kinetics of RomR relocalisation therefore introduce a refractory or relaxation period during which it is not possible to effect a reversal until sufficient RomR has accumulated. However, the above mechanism is critically incomplete to provoke the switch, since FrzX is also required as a trigger.

To test whether a combined mechanism involving FrzX can explain the reversal switch, we added the dynamics of FrzX~P to our earlier mathematical model. A simple and thermodynamically consistent mode of action of FrzX~P is to mediate the inhibitory effect proposed in our earlier model of MglA-GTP on the polar binding of MglB (Figure 2A and Supplementary Information). Based on our experimental data, we assume that FrzX~P is recruited to the pole by MglB (Figure 4B). The interactions between MglA, MglB, RomR and FrzX and the associated differential equations are presented in Figures 6A,B. Solving the equations led to the expected properties: in WT cells, the re-localisation of RomR introduced a refractory period and primed the reversal event, which was eventually provoked by the action of FrzX~P (Figure 6C). The polarity module can therefore be decomposed into two components: the MglA/MglB/RomR relaxation oscillator, with these oscillations “gated” by a FrzX~P trigger (Figure 6A), together forming a gated relaxation oscillator.

**Figure 6.**
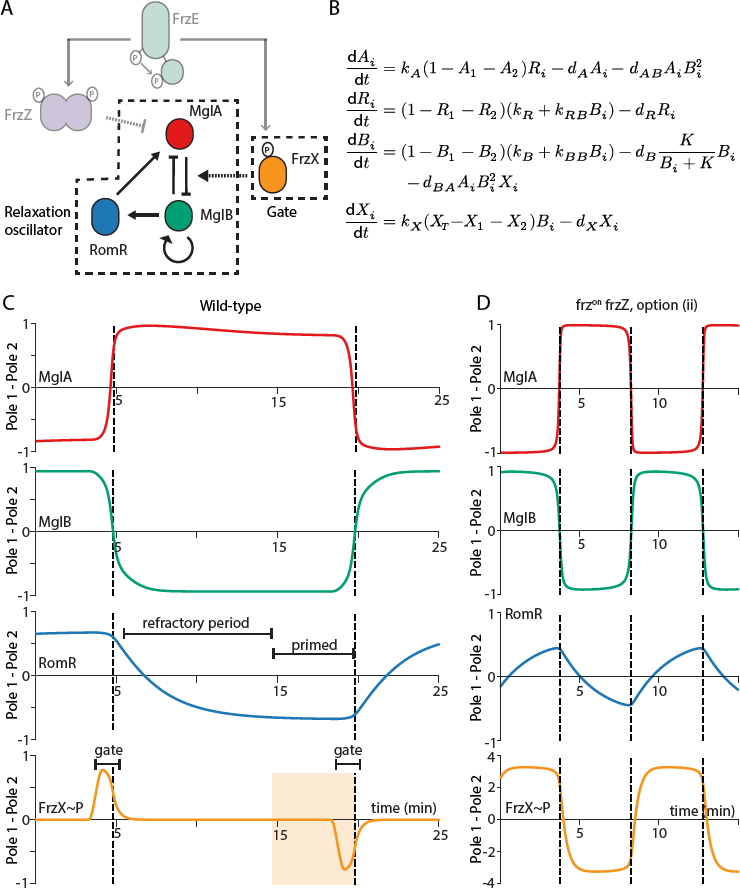
A gated relaxation oscillator model of the polarity switching mechanism captures most of the experimental data. **(A)** The polarity module of Figure 2A with the addition of FrzZ and FrzX. The kinase FrzE phosphorylates FrzZ and FrzX. The former is hypothesised to promote the unbinding of MglA, while the latter is required for MglA(-GTP) to promote unbinding of MglB. Dashed lines indicate the relaxation oscillator and gate components, respectively. Arrows representing recruitment of FrzZ~P and FrzX~P are not shown for clarity. **(B)** The equations of the refined model, updated from those in Figure 2C. X represent the relative polar copy numbers of FrzX~P. Phosphorylation of FrzX is incorporated as a pulse in the levels of the phosphorylated protein (relative total copy number: X_T_). The effect of FrzZ is incorporated globally by modifying the rate *dA* (Supplementary Information). **(C-D)** Simulated profiles of MglA, MglB, RomR and FrzX~P, generated by the equations in **(B)**, showing the difference in the polar levels for **(C)** WT and **(D)** *frz^on^ frzZ* cells (using option (ii)). Phosphorylation pulses in **(C)** have width 1.3 min. Pulses induce a switch if they occur after sufficient RomR has accumulated at the lagging pole (indicated by the shaded region), which primes it for the next switch, and not during the earlier the refractory period. Phosphorylation of FrzX~P is continuous in **(D)**. Dashed lines are as in Figure 2D. See Supplementary Information for parameter values.

We next simulated a *frz^on^* background, with constitutively high levels of FrzX~P (Supplementary Information) (Figure S6A). Switching is now more rapid, with reversals happening on a timescale faster than the intrinsic RomR dynamics (Figure 6C), leading to only weak RomR oscillations. Due to the high levels of FrzX~P, the required threshold of RomR is reduced so much that, immediately after a reversal, the new lagging pole already has sufficient RomR (and thus MglA) bound for another reversal. Since reversals occur with approximately symmetrical RomR levels at both poles (Figure S6A), consistent with our experimental observation (Figure 5G), the trigger function of FrzX~P is then clearly revealed (Figure S6A). In a *frz^on^* background, the system is still effectively a relaxation oscillator, but where now FrzX~P rather than RomR acts as the slow component. To further confirm that dynamical changes in RomR levels are inessential, we simulated artificially constant and symmetric RomR in the *frz^on^* background (Figure S6B). As expected, reversals still occurred, demonstrating the key trigger role of FrzX~P.

We next used our simulations to investigate the effect of deleting *frzZ*. Given that FrzZ~P localises to the leading pole and promotes reversals, we considered two possibilities for its mode of action: (i) FrzZ~P promotes the polar switching of MglA by accelerating the re-localisation of RomR, or (ii) FrzZ~P favors the unbinding of MglA from the leading pole without affecting the dynamics of RomR, by, for example, dissociating MglA from RomR. These two options were incorporated into the model implicitly by alterations in the kinetic parameters. We found that option (i) is likely incorrect, because removing FrzZ from the model in the *frz^on^* background (and thus simulating a *frz*^on^ *frzZ* background) resulted in a more symmetric RomR distribution at the poles (compare Figure S6A,C), whilst we observed the opposite experimentally (compare Figure 5C and 5F). We therefore pursued option (ii) (Figure 6A).

In a simulated *frz^on^ frzZ* background, using option (ii), we found that the reversal switch now became dependent on RomR accumulation to a sufficiently high level at the lagging pole (Figure 6D), as observed experimentally (Figure 5C,D). In this case, the FrzX~P trigger is quickly primed, but with slower MglA dynamics, RomR must now accumulate to high levels to recruit enough MglA to provoke the switch, similar to our original relaxation oscillator model. This reasoning was confirmed when we again simulated artificially constant and symmetric RomR: unlike in the *frz^on^* background simulated above, no reversals were seen, confirming the key role of RomR dynamics (Figure S6D). The resulting increase in RomR asymmetry in a *frz^on^* background due to the removal of FrzZ seen in our simulations (compare Figure 6D and Figure S6A) was also consistent with our experimental observations (compare Figure 5C and Figure 5F).

Finally, we found that in the absence of FrzZ alone, and thus slower MglA dynamics than the wild-type, simulated reversals were only obtained with sustained FrzX activity (Figure S6E). This result was consistent with previous observations and the restoring effect of IAA to the *frzZ* mutant^4^ (Figure 3E, Figure S5B,C).

Overall, the modified model has rationalized our findings on RomR and FrzX: the system behaves like a gated relaxation oscillator with FrzX acting as the gate or trigger. The relaxation property means that stimulation by FrzX~P is only effective if sufficient time has passed since the previous switch to allow RomR to prime the lagging pole, by recruiting MglA at a sufficiently high rate. However, in contrast to the standard gated oscillators in electronics^21^, which return to a default state, the system here remains in its current state after removal of the input (FrzX~P). Furthermore, both the experimental data and simulations suggest that FrzZ~P increases the MglA off-rate, thus decreasing the threshold of RomR required to prime the pole for a reversal. This effect shortens the RomR refractory period, allowing for more frequent reversals.

## Discussion

In this work, we have systematically dissected the dynamics of the *Myxococcus* polarity module. Using a combination of mathematical modelling and experiments, we discovered that our original three-protein relaxation oscillator model, where Frz signalling enters through modulation of the RomR dynamical timescale, was too simple. This result motivated our experimental search for an additional Frz response regulator protein, leading to the identification of FrzX. Subsequent characterisation of the role of FrzX, then led to an improved model in which FrzX acts as the trigger of a gated relaxation oscillator. This model successfully accounts for many features of the polarity switching dynamics in *Myxococcus*.

Controlling reversals in this way combines many of the advantages of both a switch and an oscillator. The presence of a relaxation oscillator naturally causes the polarity apparatus to reverse poles, an essential feature. However, direct control of the relaxation oscillator period would require the tuning of a continuous variable (the RomR timescale), a nontrivial task if a wide response range is to be obtained. Employing a gating mechanism bypasses these constraints and only requires that the levels of the input (FrzX~P) exceed a given threshold when a reversal is required and stay below it otherwise. Furthermore, the presence of the refractory period may also be advantageous. Unlike swimming bacteria, which can change their direction of movement in 3D (usually via tumbling), *Myxococcus* has a binary choice: reverse or not. A refractory period can ensure that a cell cannot be stimulated again immediately after it reversed which could be important for cooperative motility behaviors. Indeed, mathematical models have predicted that, in addition to an internal biochemical clock, a refractory period is required for rippling behaviour^6,27–29^. We have shown that this refractory period is shortened when Frz signalling is high, but is lengthened by the absence of FrzZ. Such fine-tuned control is clearly important because the *frzZ* mutant is defective in most developmental processes.

Another necessary consequence of our gated switch mechanism is that the system becomes sensitive to variations in the level of FrzX~P. Assuming that the pool of FrzX does not vary abruptly between cells, these fluctuations could be introduced not only by varying levels of FrzE activity but also due to the existence of a high auto-phosphatase activity of FrzX, as suggested by our inability to capture FrzX~P on SDS-PAGE gels (Figure 3D). These effects could be important because wild-type *Myxococcus* cells show a broad distribution of reversal frequencies^4^, but all mutants with narrow distributions, be they in the fast *(frz^on^)* or slow *(frzZ)* ranges, are severely impaired in development^30^.

We have found that RomR acts as an oscillating scaffold protein, regulating the polar concentration of MglA. Our data indicates that RomR is never fully localised to the lagging cell pole, which likely explains why MglA remains attached to the leading cell pole even if the majority of RomR localises to the opposite pole. We have further proposed that FrzX~P could be responsible for triggering the inhibitory effect of MglA-GTP on MglB (see Supplementary Information for further discussion). Note, however, that FrzX alone cannot be the long sought after MglA Guanine nucleotide Exchange Factor (GEF) because MglA is active in absence of FrzX. Moreover, we cannot of course exclude a more complex sequence of triggering events occurring downstream from FrzX, perhaps involving as yet unidentified polar proteins.

In conclusion, the *Myxococcus* polarity system forms a new type of genetic circuit in which two polar response regulators mediate two distinct controls. One (RomR) ramps up slowly as part of a relaxation oscillator and must exceed a critical level, while the other (FrzX~P) then acts as a critical checkpoint or gate to trigger a reversal. This flexible gated relaxation oscillator architecture, incorporating a checkpoint into an oscillator, allows a wide range of reversal dynamics in response to environmental signals. It is therefore likely that this type of regulation also occurs in other rhythmic biological systems.

## Methods

### Bacterial strains, growth conditions and genetic constructs

Strains, plasmids and primers used for this study are listed in Tables S1, S2 and S3. All genetic mutants were constructed in the *Myxococcus xanthus* DZ2 strain^30^. *M. xanthus* DZ2 was grown at 32°C in CYE rich media, as previously described^30^. Plasmids were introduced in *M. xanthus* by electroporation. Mutants and transformants were obtained by homologous recombination based on a previously reported method^30^. Complementation, expression of the fusion and mutant proteins were either obtained by ectopic integration of the genes of interest at the Mx8-phage attachment site^12^ under the control of their own promoter in appropriate deletion backgrounds, or by expression from the endogenous locus (Table S2). Clean replacement and deletions were constructed as previously reported.

The plasmid to replace *frzX* contains an insert encompassing 850 bp upstream from the *frzX* coding sequence to 850 bp immediately downstream of the *frzX* coding sequence and synthesized into the pBluescriptII SK(+) plasmid by Biomatik. This plasmid was then digested by HindIII/EcoRI restriction enzymes and the insert was ligated into the pBJ114 vector.

For clean replacement of the *romR* gene by *romR-sfgfp* at the endogenous locus, pBJ114-*romR-sfgfp* was constructed by amplifying and fusing 850 bp upstream from the *romR* stop codon, the *sf-gfp* gene and 850 bp immediately downstream from the *romR* coding sequence, by overlap PCR. The resulting PCR product was digested by the EcoRI*/HindIII* restriction enzymes and ligated into the pBJ114 vector. The insert was verified by sequencing.

The plasmids carrying the *romR^D53E/N^* point mutations were constructed by PCR with oligonucleotides carrying the mutation amplifying a fragment encompassing 500 bp upstream from the mutation site with a fragment encompassing 1000 bp downstream from the point mutation. The two fragments were fused by overlap PCR, digested by the *HindIII/XbaI* restriction enzymes and cloned into the pBJ114 plasmid. The insert was verified by sequencing.

For replacement of the *frzZ* gene by *frzZ-mcherry* at the locus, pBJ1*14-frzZ-mcherry* was constructed by amplifying independently 857 bp encompassing the 5’ end of *frzZ* and the *mcherry* gene. The two fragments were fused by overlap PCR. The resulting PCR product was digested by the BamHI/EcoRI restriction enzymes and ligated into the pBJ114 vector. The insert was verified by sequencing.

Complementation of the *frzX* deletion was obtained by expressing *frzX* under the control of its own promoter from the pSWU30 plasmid, an integrating plasmid that recombines at the Mx8 site. A fragment encompassing *frzX* and the upstream 200 bp sequence containing the promoter was digested using the EcoRI/BamHI restriction enzymes and ligated into the pSWU30 vector. *pSWU19-frzX^D55A^* was constructed with the same external primers as the *frzX* complementation, but with internal overlapping primers carrying the point mutation. The fragments were fused using overlap PCR. The resulting fragment was digested and ligated into pSWU19, a pSWU30 derivative containing a Kanamycin resistance cassette^12^.

GFP-FrzX was expressed in a *frzX* deletion background by expressing *sfGFP-frzX* from the *frzX* promoter integrated at the Mx8 site. For this, the *frzX* promoter region upstream was fused upstream from the *sf-gfp* gene, itself fused in-frame with the *frzX* gene. The resulting PCR product was digested using the BamHI/EcoRI restriction enzyme and ligated into the pSWU19 vector.

### Phenotypic assays

Development assays were performed as previously described^30^. Cells were grown up to exponential phase and concentrated at Optical Density OD = 5 in TPM buffer (10 mM Tris-HCl, pH 7.6, 8 mM MgSO4 and 1 mM KH2PO4). Then they were spotted (10 μL) on CF 1.5% agar plate. Colonies were photographed after 72 h of incubation at 32°C. Developmental assays in the presence of Isoamyl alcohol (IAA, Sigma Aldrich) were performed similarly, except that plates also contained IAA at appropriate concentrations.

### Cloning, expression and purification of *M. xanthus* Frz system proteins

The cloning, expression and purification of *M. xanthus* Frz system proteins were performed as previously described^4^. Briefly, the genes encoding FrzZ, FrzX, RomR and RomR^RR^ were amplified by PCR using *M. xanthus* DZ2 chromosomal DNA as a template, with the forward and reverse primers listed in Table S3. The amplified product was digested with the appropriate restriction enzymes, and ligated into either pETPhos or pGEX. All constructs were verified by DNA sequencing. The generated plasmids were used to transform *E. coli* BL21(DE3)Star cells in order to overexpress His-tagged or GST-tagged proteins. Recombinant strains harboring the different constructs were used to inoculate 400 ml of LB medium supplemented with glucose (1 mg/mL) and ampicillin (100 μg/ml). The resulting cultures were incubated at 25°C with shaking until the optical density of the culture reached an OD = 0.6. IPTG (0.5 mM final) was added to induce overexpression, and growth was continued for 3 extra hours at 25°C. Purification of the His-tagged/GST-tagged recombinant proteins was performed as described by the manufacturer (Clontech/GE Healthcare).

### *In vitro* autophosphorylation assay

The *in vitro* phosphorylation assay was performed as described^4,15^, with *E. coli* purified recombinant proteins. Briefly, 4 μg of FrzE^kinase^ was incubated with 1 μg of FrzA and increasing concentrations (0.5 to 7 μg) of FrzCD in 25 μl of buffer P (50 mM Tris-HCl, pH 7.5; 1 mM DTT; 5 mM MgCl_2_; 50mM KCl; 5 mM EDTA; 50μM ATP, 10% glycerol) supplemented with 200 μCi ml^-1^ (65 nM) of [γ-33P]ATP (PerkinElmer, 3000 Ci mmol^-1^) for 10 minutes at room temperature in order to obtain the optimal FrzE^kinase^ autophosphorylation activity. Each reaction mixture was stopped by addition of 5 × Laemmli and quickly loaded onto SDS-PAGE gel. After electrophoresis, proteins were revealed using Coomassie Brilliant Blue before gel drying. Radioactive proteins were visualized by autoradiography using direct exposure to film (Carestream).

### Fluorescence microscopy

Time-lapse experiments were performed as previously described^4^, using an automated and inverted epifluorescence microscope TE2000-E-PFS (Nikon, France), with a 100×/1.4 DLL objective and a CoolSNAP HQ2 camera (Photometrics). The microscope is equipped with “The Perfect Focus System” (PFS) that automatically maintains focus so that the point of interest within a specimen is always kept in sharp focus at all times, in spite of any mechanical or thermal perturbations. Images were recorded with Metamorph software (Molecular Devices). All fluorescence images were acquired with a minimal exposure time to minimize bleaching and phototoxicity effects.

### FRAP experiments

Image acquisition and FRAP measurements were performed on a custom-made upright monolithic aluminum microscope^31^ with a 100 × /1.49 N.A. objective (Nikon), iXon DU-897 cooled EMCCD camera (Andor Technology), and a homemade LabView software package (National Instruments).

Photobleaching was achieved by focusing an argon ion laser to a diffraction-limited spot on the specimen for a pulse of ≈5 s. Wide-field fluorescent images of the cells were acquired before and after photobleaching by custom time-lapse recording of digital images with 16-bit grey levels. Images of both wild-type MglA-YFP cells and MglB-YFP cells were obtained every 0.4 s for 30 s. Images of wild-type RomR-GFP cells were obtained every 10 s for 4 min.

### Bioinformatics

Genomic contexts: close FrzX homologs identified by BLAST were analysed using Microbial Genomic Context Viewer (http://mgcv.cmbi.ru.nl/). Protein sequence alignment was performed using Mafft (https://toolkit.tuebingen.mpg.de/mafft) and Ali2D for secondary structure prediction (https://toolkit.tuebingen.mpg.de/ali2d).

### Mathematical modelling

The model is described by a set of ordinary differential equations. The equations were solved numerically using the *ode45* solver in Matlab (The Mathworks Inc.). We mimicked signalling from the Frz system to the response regulator FrzX using a square wave smoothened with the *smooth* function. The bifurcation diagram was created using the matcont toolbox (https://matcont.sourceforge.io). See main text and Supplementary Information for detailed description and justification.

## Quantification and Statistical Analysis

### Reversal scoring assays

Reversals were scored using previously described microfluidic single cell carboxymethylcellulose based assays, allowing modulation of Frz-signaling intensity with IAA^4^. In this assay, cells are moving using the S-motility system and reversals are scored in the presence of IAA (0.1% or 0.15%) or without IAA (for mutants with a *frz^on^* mutation). For this, homemade PDMS glass microfluidic chambers^32^ were treated with 0.015% carboxymethylcellulose after extensive washing of the glass slide with water. For each experiment, 1 mL of a CYE grown culture of OD = 0.5–1 was injected directly into the chamber and the cells were allowed to settle for 5 min. Motility was assayed after the chamber was washed with TPM 1mM CaCl_2_ buffer^4^. IAA solutions made in TPM 1mM CaCl2 buffer at appropriate concentrations (0.1% or 0.15%) were injected directly into the channels. Reversals were scored using two different analyses, since rapid directional changes unlinked to reversals but linked to motility engine activity (so-called stick-slip motions^4,33^) can occur infrequently accounting for a low number of false positives. In mutants with no stick-slip motions, directional changes were monitored in contrast images acquired every 15 s for 30 min (see below) with 0.15% IAA, or without for mutants carrying the *frz^on^* mutation. To discriminate stick-slip motions from *bona fide* reversals in certain mutants, oscillations of FrzS-YFP proteins were scored instead of directional movements in the presence of 0.1% IAA (see below). This method has proven particularly accurate in determining the effect of a given gene in reversal control^4^. Here, we use this method to unambiguously characterise the role of FrzX.

### Cell tracking

Image analysis was performed with a specific library of functions written in Python and adapted from available plugins in FIJI/ImageJ^34^. Cells were detected by thresholding the phase contrast images after stabilization. Cell were tracked by calculating all object distances between two consecutive frames and selecting the nearest objects. The computed trajectories were systematically verified manually and, when errors were encountered, the trajectories were removed. The analysis of the trajectories was performed using a Python script that calculates the angle formed by the line segments between the center of the cell at time *t*, the center at time *t*-1 and the center at time *t*-1.

Directional changes were scored as reversals when cells switched their direction of movement, and the angle between segments was less than 90°. For non-reversing strains, the reversal frequency was calculated by dividing the number of directional changes by the number of tracked frames, and, since images were acquired every 15s, this number was multiplied by 240 to generate a reversal frequency per hour and plotted. For strains that frequently reversed, the mean time between two reversals for each cell was plotted. Plotting was performed using the software R (http://www.R-project.org/).

To further discriminate *bona fide* reversal events from stick-slip motions, the fluorescence intensity of FrzS-YFP was measured at cell poles over time. For each cell that was tracked, the fluorescence intensity and reversal profiles were correlated to distinguish bona fide reversals from stick-slip events with the R software. When a directional change was not correlated to a switch in fluorescence intensity, this change was discarded as a stick-slip event. The reversal frequency was then calculated by dividing the number of bona fide reversals by the number of tracked frames, and, since images were acquired every 15s, this number was multiplied by 240 to generate a reversal frequency per hour and plotted.

Statistics were performed using R: the Wilcoxon test was used when the number of cells was less than 40 in at least one of the two populations compared, while the student test (t-test) was used when the numbers of cells was higher than 40.

### Generating single cell traces

Tracking of fluorescent polar clusters was performed manually with the MTrackJ plugin under FIJI after background subtraction and using a local cursor snapping to detect the maximum of fluorescence intensity at a pole. Normalized fluorescence intensities were calculated by normalizing the difference in polar signal by its maximum absolute value: I=(Pole1-Pole2)/max(abs(Pole 1-Pole 2)).

### Switching timescales

Measurement of the timescale of polar switching, i.e. how long it takes a fluorescent foci to switch poles, was performed using an automated method under the FIJI/ImageJ plugin MicrobeJ^35^. Cell outlines and tracks were obtained based on phase contrast images and used to extract the subcellular fluorescence (using the ‘profile’ option). In this methodology, MicrobeJ performs a projection of the fluorescent signal onto the medial axis of the cell, returning the projected signal at discrete points along this axis (beginning and ending at each pole, and approximately 1 pixel apart). These discrete points define the boundary of transverse segments along the medial axis. The R script determines, for each cell on each frame, the area A(i) and the total fluorescent signal S(i) of each segment i. The sum of these values are the cross-sectional cell area (A_cell) and the total fluorescence of the cell respectively. The mean fluorescence *intensity* of the cell (Int_cell) is the latter divided by the former. The polar regions were defined to be the first and last 5 segments of the cell and the corresponding mean polar fluorescence intensities, P1 and P2, are given by dividing the sum of the fluorescent signal, S(i), of these segments by the sum of the corresponding segment areas A(i). The mean cytoplasmic (CYTO) fluorescence intensity was defined as the summed intensity of the remaining segments divided by the sum of the corresponding segment areas. A correction for background fluorescence was made by subtracting the background intensity (extracted from the MicrobeJ output). A photobleaching correction was also added by normalizing to the whole cell intensity (Int_cell). We removed the effect of autofluorescence and diffuse cytoplasmic signal by subtracting CYTO from P1 and P2. We denote these corrected mean polar intensities by Y1 and Y2, respectively.

We found that working with the difference between the two polar intensities was less noisy than looking at the poles individually and we define the normalised difference as Y=(Y1-Y2)/(Y1+Y2). The profile, Y was then smoothened using a Savitzky-Golay filter (Fig. S1D, upper plot). Nevertheless, variation in Y made it difficult to say when precisely a switch began and ended in all cases. However, one time point that can be determined robustly is the time of maximum (or minimum) slope, i.e. the time at which the signal changes fastest. We therefore used this measure as the basis of our approach. To calculate the slope maxima and minima, we used a Savitzky-Golay derivative filter to calculate a smoothened derivative directly from the unsmoothed profile (Fig. S1D, lower plot). We also take the time period between consecutive minima/maxima to define individual cycles (and the cycle duration). This generally agrees with the times of parity in the signal between the poles. We define the switching timescale to be the duration of the switch if the signal were to change continuously at its maximum measured rate. This is calculated as twice the maximum absolute value of Y within a cycle divided by the absolute value of the maxima/minima of the derivative (Fig. S1D). Note that each cycle has two associated switching timescales.

### Correlating fluorescence intensities and reversals

To correlate fluorescence intensities and reversals, the time interval between a reversal, detected by the first time frame where a directional change can be measured, and the maximum of fluorescence intensity of a given GFP/mCherry fusion was scored for each reversal event (n). Negative time values were given if the maximum intensity was measured before a reversal and positive time values were given if the maximum intensity was measured after a reversal. A zero value means that a fluorescence maximum is observed exactly at the reversal detection time. Note that due to phototoxicity and photobleaching, the time resolution of the measurements is 30 s and thus detection of the exact reversal time has a resolution equal to this interval.

### FRAP analysis

The analysis of FRAP data was performed using a custom MATLAB script. Data stored as multiplane tiff images were read in and presented to the user to select a region of interest as well as the bleached region. After background subtraction, the mean intensity within the region of interest was corrected for acquisition bleaching by multiply the intensities by an exponential exp(kt), the coefficient, k, having being obtained by fitting to the acquisition bleaching curve of another cell. Finally, we normalized to a prebleach intensity of 1. The resulting curves were fit using non-linear least squares to the equation f(t)=a*(1-exp(-b*t))+c, where *a* is a measure of the amplitude of the recovery, *b* is the inverse of the timescale of recovery and *c* is the fraction of the initial intensity that was not bleached.

Other than the intensity in the bright focus near the pole, the background intensity in the cell is uniform along the long axis of the cell, even at the fastest timescales that we investigated (2.5 frames/second). As described in the Supporting Information section describing the three compartment model, this implies that the timescale of FRAP recovery is dominated not by the diffusion of free protein through the cytoplasm, but the binding and unbinding kinetics. Following the work of Sprague et al.^36^, we interpret the constant *b* from each cell to be an estimate of *k_off_*. The half-time of recovery, t_1/2_, is ln(2)/*b*.

Statistics were performed using R: the Wilcoxon test was used when the number of cells (n) was less than 40 in at least one of the two populations compared, and the student test (t-test) was used for a number of cells higher than 40.

## Acknowledgments

We wish to thank Lotte Sogaard-Andersen and Ulrich Gerland for discussions and Victor Sourjik for comments on the manuscript. MG was the recipient of an ARC fellowship (DOC20140601482). EM is the recipient of an AMIDEX “Académie d’excellence” thesis fellowship (n°ANR-11-IDEX-0001-02). SL is funded by an ANR program “BACTOCOMPASS” TM is the recipient of an ERC starting grant “DOME 261105” and an ANR program “BACTOCOMPASS”. RV was supported by the French Infrastructure for Integrated Structural Biology (FRISBI) ANR-10-INBS-05.

## Author contributions

Conceptualization, MG, SMM, EM, SL, MH and TM; Methodology, MG, SMM, EM, SL, VM, MH and TM.; Investigation, MG, SMM, EM, SL, LM, GB and YZ; Writing - Original Draft, MH and TM; Writing - Review & Editing, MG, SMM and EM; Visualization: MG, SMM, EM, SL, LE, MH and TM; Funding Acquisition, MH and TM; Resources, BB, RV, JS and MV; Mathematical analysis: SMM and MH; Supervision, MH and TM.

## Supplemental figure titles and legends

### Figure S1: Recovery times of polarity proteins MglA-YFP, MglB-YFP and RomR-GFP. Related to Figure 1.

**(A)** A schematic of the switching timescale measurement. The normalised difference between polar signals, Y, is determined from time-lapse analysis (upper plot, blue dots). Savitzky-Golay filters are used to obtain a smoothed profile (upper plot, red line) and a smoothed profile of the derivative (lower plot, blue line). The duration of a switching cycle is defined to be the time between a peak and subsequent (or preceding) valley in the derivative profile (indicated for the first cycle by shading and crosses). Since the derivative is simply the angle between the tangent (black dashed lines) to the profile and the x-axis (blue triangles), we can use values of the maxima/minima of the derivative to define appropriate timescales: the two switching timescales associated with each cycle are defined to be twice the maximum absolute value, M, of (smoothened) Y within each cycle divided by the absolute value of the minimum/maximum of the derivative profile (S).

**(B)** Top: Example of single-cell recovery after photobleaching for each polarity protein. The fluorescence intensity dynamics in the bleached regions (blue crosses and lines), corrected for bleaching, were fit to exponential curves (red lines) by non-linear least squares fitting. Bottom: Timelapse images (0.4 s time frame for MglA-YFP and MglB-YFP, 10 s time frame for RomR-GFP) of single-cell recovery after photobleaching.

**(C)** MglA-YFP and **(D)** MglB-YFP recovery times as determined by FRAP in WT, *frz^on^* and *frzE* mutants. The lower and upper boundaries of the boxes correspond to the 25% and 75% percentiles, respectively. The median is shown as a red line and the whiskers represent the 10% and 90% percentiles.

### Figure S2: RomR is not phosphorylated by FrzE. Related to Figure 3.

**(A)** Schematic of RomR domain structure.

**(B)** RomR^RR^ is not phosphorylated by the FrzE kinase *in vitro*. Autoradiogram of P^32^-labelled FrzE in the presence of RomR^RR^.

**(C)** Developmental phenotypic assays showing the absence of fruiting body formation in a *romR* mutant. *romR* is essential for fruiting body formation but point mutations at the phosphorylatable aspartate of the RomR response regulator domain do not affect fruiting body formation. Scale bar: 1 mm.

### Figure S3: FrzX is a target of FrzE and is essential for reversals. Related to Figure 3.

**(A)** Genomic context of genes encoding FrzX and FrzX-like proteins in the *Myxococcales*and *Anaeromyxobacter* species.

**(B)** and **(C)** Fruiting body formation of **(B)** WT, *frzX* mutant and *frzX* mutant complemented with an ectopic copy of the *frzX* gene, (C) an ectopic copy of the *fizX^D55A^* gene and an ectopic copy of the *gfp-frzX* gene. Scale bar: 1 mm.

### Figure S4: Localization dependencies between the *frz* proteins and the polarity proteins. Related to Figure 4

**(A)** Images showing localization of MglA-YFP, MglB-YFP, RomR-GFP in the *frzX* mutant. Scale bar = 2 μm.

**(B)** Images showing localization of FrzZ-GFP in the WT and *frzE* and *frzX* mutants. Scale bar = 2 μm.

### Figure S5: Dynamics of GFP-FrzX and RomR-GFP in a *frz^on^* background and a *frzZ* background in the presence of IAA. Related to Figure 5.

**(A)** Top: time lapse images of GFP-FrzX dynamics from a single reversing cell in a *frz^on^* background (30 s time frames). Bottom: difference in intensity of GFP-FrzX between the two poles normalized to maximum absolute value of the difference. R: Reversals, detected by a clear directional change of the cell. Scale bar = 2 μm.

**(B),(C)** and **(D)** Same as **(A)** but for **(B)** GFP-FrzX with 0.1% IAA in a *frzZ* background (15 s time frames), **(C)** RomR-GFP with 0.1% IAA in a *frzZ* background and **(D)** GFP-FrzX (green trace) and RomR-mCh (red trace) in a WT background with 0.3% IAA (30 s time frames). Scale bar = 2 μm

### Figure S6: Simulated polarity protein dynamics. Related to Figure 6.

**(A)** Simulated profiles of MglA, MglB, RomR and FrzX~P, generated by the equations in Figure 6B, showing the difference in polar levels in the *frz^on^* background. Phosphorylation of FrzX~P is continuous. For all panels, dashed lines are as in Figure 2D. See Supplementary Information for parameter values.

**(B)** Simulated profiles in *frz^on^* cells as in (A) but with RomR levels held constant and symmetric. Asymmetry in polar RomR is not required for oscillations.

**(C)** Simulated profiles in *frz^on^ frzZ* cells, assuming FrzZ promotes RomR unbinding (option (i) of main text). Parameters are same as in (A) but with a one order of magnitude lower rate of RomR unbinding.

**(D)** Simulated profiles in *frz^0n^ frzZ* cells (with option (ii) of main text) as in Figure 6D, but with RomR levels held constant and symmetric. In this case, asymmetry in polar RomR is clearly required for oscillations.

**(E)** Simulated profiles in *frzZ* cells (option (ii) of main text). Shown is the effect of phosphorylation pulses of differing durations (1.3, 2 and 3 mins). Unlike the wild-type, a 1.3 min pulse is not sufficient to induce a switch, nor is a 2 min pulse. However, a 3 min pulse (or greater) is sufficient and induces a polar switch if it occurs when sufficient RomR has accumulated at the lagging pole (indicated by the shaded region). See Supplementary Information for parameter values and justification for modelling FrzX activity as pulsatile.

## Supplemental tables

**Table S1. Bacterial strains used in this study. Related to STAR Methods**

**Table S2. Plasmids used in this study. Related to STAR Methods**

**Table S3. Primers used in this study. Related to STAR Methods**

## References

1. Goldbeter, A., Gérard, C., Gonze, D., Leloup, J. C. & Dupont, G. Systems biology of cellular rhythms. in FEBS Letters 586, 2955–2965 (2012).

2. Cohen, S. E. & Golden, S. S. Circadian Rhythms in Cyanobacteria. Microbiol. Mol. Biol. Rev. 79, 373–385 (2015).

3. Howard, M. & Kruse, K. Cellular organization by self-organization: Mechanisms and models for Min protein dynamics. Journal of Cell Biology 168, 533–536 (2005).

4. Guzzo, M. et al. Evolution and Design Governing Signal Precision and Amplification in a Bacterial Chemosensory Pathway. PLOS Genet. 11, e1005460 (2015).

5. Sliusarenko, O., Neu, J., Zusman, D. R. & Oster, G. Accordion waves in Myxococcus xanthus. Proc. Natl. Acad. Sci. 103, 1534–1539 (2006).

6. Igoshin, O. A., Goldbeter, A., Kaiser, D. & Oster, G. A biochemical oscillator explains several aspects of Myxococcus xanthus behavior during development. Proc. Natl. Acad. Sci. U. S. A. 101, 15760–5 (2004).

7. Chang, Y.-W. et al. Architecture of the type IVa pilus machine. Science (80-.). 351, aad2001–aad2001 (2016).

8. Faure, L. M. et al. The mechanism of force transmission at bacterial focal adhesion complexes. Nature 539, 530–535 (2016).

9. Leonardy, S. et al. Regulation of dynamic polarity switching in bacteria by a Ras-like G-protein and its cognate GAP. EMBO J. 29, 2276–2289 (2010).

10. Mauriello, E. M. F. et al. Bacterial motility complexes require the actin-like protein, MreB and the Ras homologue, MglA. EMBO J. 29, (2009).

11. Treuner-Lange, A. et al. The small G-protein MglA connects to the MreB actin cytoskeleton at bacterial focal adhesions. J. Cell Biol. 210, 243–56 (2015).

12. Zhang, Y., Franco, M., Ducret, A. & Mignot, T. A bacterial Ras-like small GTP-binding protein and its cognate GAP establish a dynamic spatial polarity axis to control directed motility. PLoS Biol. 8, e1000430 (2010).

13. Miertzschke, M. et al. Structural analysis of the Ras-like G protein MglA and its cognate GAP MglB and implications for bacterial polarity. EMBO J. 30, 4185–4197 (2011).

14. Kaimer, C. & Zusman, D. R. Regulation of cell reversal frequency in Myxococcus xanthus requires the balanced activity of CheY-like domains in FrzE and FrzZ. Mol. Microbiol. 100, 379–95 (2016).

15. Inclán, Y. F., Vlamakis, H. C. & Zusman, D. R. FrzZ, a dual CheY-like response regulator, functions as an output for the Frz chemosensory pathway of Myxococcus xanthus. Mol. Microbiol. 65, 90–102 (2007).

16. Inclán, Y. F., Laurent, S. & Zusman, D. R. The receiver domain of FrzE, a CheA-CheY fusion protein, regulates the CheA histidine kinase activity and downstream signalling to the A-and S-motility systems of Myxococcus xanthus. Mol. Microbiol. 68, 1328–1339 (2008).

17. Kaimer, C. & Zusman, D. R. Phosphorylation-dependent localization of the response regulator FrzZ signals cell reversals in Myxococcus xanthus. Mol. Microbiol. 88, 740–753 (2013).

18. Leonardy, S., Freymark, G., Hebener, S., Ellehauge, E. & Søgaard-Andersen, L. Coupling of protein localization and cell movements by a dynamically localized response regulator in Myxococcus xanthus. EMBO J. 26, 4433–4444 (2007).

19. Keilberg, D., Wuichet, K., Drescher, F. & Søgaard-Andersen, L. A Response Regulator Interfaces between the Frz Chemosensory System and the MglA/MglB GTPase/GAP Module to Regulate Polarity in Myxococcus xanthus. PLoS Genet. 8, (2012).

20. Zhang, Y., Guzzo, M., Ducret, A., Li, Y.-Z. & Mignot, T. A dynamic response regulator protein modulates G-protein-dependent polarity in the bacterium Myxococcus xanthus. PLoS Genet. 8, e1002872 (2012).

21. Barnaal, D. Analog Electronics for Scientific Application. (Waveland Press, 1989).

22. Bowman, G. R. et al. Oligomerization and higher-order assembly contribute to sub-cellular localization of a bacterial scaffold. Mol. Microbiol. 90, 776–795 (2013).

23. Laloux, G. & Jacobs-Wagner, C. Spatiotemporal control of PopZ localization through cell cycle-coupled multimerization. J. Cell Biol. 201, 827–841 (2013).

24. Skerker, J. M., Prasol, M. S., Perchuk, B. S., Biondi, E. G. & Laub, M. T. Two-component signal transduction pathways regulating growth and cell cycle progression in a bacterium: a system-level analysis. PLoS Biol. 3, e334 (2005).

25. Thomas, S. A., Brewster, J. A. & Bourret, R. B. Two variable active site residues modulate response regulator phosphoryl group stability. Mol. Microbiol. 69, 453–465 (2008).

26. McBride, M. J., Weinberg, R. a & Zusman, D. R. ‘Frizzy’ aggregation genes of the gliding bacterium Myxococcus xanthus show sequence similarities to the chemotaxis genes of enteric bacteria. Proc. Natl. Acad. Sci. U. S. A. 86, 424–8 (1989).

27. Igoshin, O. A., Mogilner, A., Welch, R. D., Kaiser, D. & Oster, G. Pattern formation and traveling waves in myxobacteria: theory and modeling. Proc. Natl. Acad. Sci. U. S. A. 98, 14913–8 (2001).

28. Igoshin, O. A., Welch, R., Kaiser, D. & Oster, G. Waves and aggregation patterns in myxobacteria. Proc. Natl. Acad. Sci. U. S. A. 101, 4256–4261 (2004).

29. Zhang, H. et al. The mechanistic basis of Myxococcus xanthus rippling behavior and its physiological role during predation. PLoS Comput. Biol. 8, e1002715 (2012).

30. Bustamante, V. H., Martínez-Flores, I., Vlamakis, H. C. & Zusman, D. R. Analysis of the Frz signal transduction system of Myxococcus xanthus shows the importance of the conserved C-terminal region of the cytoplasmic chemoreceptor FrzCD in sensing signals. Mol. Microbiol. 53, 1501–1513 (2004).

31. Morgenstein, R. M. et al. RodZ links MreB to cell wall synthesis to mediate MreB rotation and robust morphogenesis. Proc. Natl. Acad. Sci. U. S. A. 112, 12510–5 (2015).

32. Ducret, A., Théodoly, O. & Mignot, T. in Bacterial Cell Surfaces 97–107 (2013). doi:10.1007/978-1-62703-245-2_6

33. Hu, W. et al. Exopolysaccharide-independent social motility of Myxococcus xanthus. PLoS One 6, (2011).

34. Schindelin, J. et al. Fiji: an open-source platform for biological-image analysis. Nat. Methods 9, 676–82 (2012).

35. Ducret, A., Quardokus, E. M. & Brun, Y. V. MicrobeJ, a tool for high throughput bacterial cell detection and quantitative analysis. Nat. Microbiol. 1, 16077 (2016).

36. Sprague, B. L. & McNally, J. G. FRAP analysis of binding: Proper and fitting. Trends in Cell Biology 15, 84–91 (2005).

